# Alpha-synuclein amyloids catalyze the degradation of ATP and other nucleotides

**DOI:** 10.1101/2025.04.03.647058

**Authors:** Claudio Castillo-Caceres, Esteban Nova, Rodrigo Diaz-Espinoza

## Abstract

Intracellular accumulation of alpha-synuclein amyloids is a main pathological hallmark in a subgroup of human neurodegenerative diseases called synucleinopathies. Cell death of energy-deprived dopaminergic neurons causes decreased dopamine levels, which underly many of the neurological symptoms in the most prevalent synucleinopathy, Parkinson’s disease. Amyloid-mediated toxicity can proceed via gain-of-function through diverse pathways. In this work, we report that alpha-synuclein amyloids can degrade adenosine triphosphate in a catalytic fashion, producing adenosine diphosphate and adenosine monophosphate. Upon prolonged incubation, all adenosine triphosphate is irreversibly consumed. Furthermore, these amyloids can also degrade all other ribonucleotides with different efficiencies, including guanosine, cytidine, and uridine triphosphates. Our findings uncover a previously unknown gain-of-function for alpha-synuclein amyloids, which may have far reaching implications for ATP and nucleotide metabolism during neurodegeneration in Parkinson’s disease and other synucleinopathies.

---

Synucleinopathies are neurodegenerative and chronic diseases affecting the human nervous system^1^. They share a common pathological feature characterized by progressive accumulation and aggregation of an intracellular protein called alpha synuclein (α−Syn) in brain cells^1,2^. Parkinson’s disease (PD) is by far the most common synucleinopathy and the second most prevalent neurodegenerative disorder in the worldwide population, only preceded by Alzheimer’s disease^3^. PD is characterized by progressive loss of motor function leading to rigidity and involuntary tremors, followed by non-motor neurological symptoms such as dementia^4,5^. The most affected brain tissue area is the substantia nigra, a midbrain zone loaded with dopaminergic neurons that fill the basal ganglia with the necessary levels of dopamine for proper control of voluntary motor^6^. Progressive death of these neurons leads to the onset of PD motor symptoms. As with most neurodegenerative disorders, to date there is no cure available for PD and other synucleinopathies.

Accumulation of abnormally folded proteins as intermolecular aggregates called amyloids is a main feature underlying most neurodegenerative diseases^7,8^. Amyloids are highly ordered and stable aggregates that can propagate and accumulate in tissues^9^. Common structural features include the formation of oligomers and fibrils that share a core and stabilizing beta sheet. In synucleinopathies, α−Syn accumulates as intracellular amyloid deposits. In PD and several other synucleinopathies, these deposits evolve into characteristic clumps called Lewy bodies^10^. Diverse evidence supports a direct association between pathology and α−Syn amyloids^11,12^. However, this association is complex and thus actively studied. Although accumulation of aggregates can partially deplete cells of α−Syn and leads to lack-of-function phenotypes, there is much evidence indicating that aggregated α−Syn species are per se toxic^13,14^. This gain-of-function can proceed through fibrillary or oligomeric states. For instance, α−Syn oligomers are highly toxic in cultured neuronal cells^15,16^. They can also easily spread and trigger synaptic impairment ^17^. Intracerebral injection of α−Syn amyloid fibrils induces parkinsonism-like symptoms in rodent animal models^18,19^. At the cellular level, α−Syn amyloids can affect the mitochondrial membrane and its components, producing oxidative stress leading to cell death^20–23^. In fact, mitochondrial dysfunction is a well-known hallmark of PD in cellular and animal models, and in human postmortem tissue^24^.

Amyloids with catalytic activity can be assembled with rationally designed small peptides, which may lead to future nanomaterials^25,26^. However, this novel feature is not restricted to synthetic non-pathological amyloids^27^. β-amyloid is an extracellular peptide released by abnormal proteolytic cleavage that accumulates as amyloid plaques in Alzheimer’s disease^28^. *In vitro*-produced β-amyloid fibrils can catalyze the hydrolysis and oxidation of several brain hormones and neurotransmitters^29^. Interestingly, amyloid fibrils assembled with recombinant α−Syn can catalyze the hydrolysis of phosphate and acetate from synthetic nitrophenyl compounds^30^. Moreover, these amyloids can modify the natural ratios of several intracellular metabolites in crude cell extracts, though no catalytic nor evidence of direct interaction was reported^31^. The same amyloids were recently shown to bind DNA and induce chemical damage^32^. These novel findings suggest that amyloid-mediated catalysis may have unforeseen and potentially key roles in pathology.

In this work, we explored whether α−Syn amyloids can directly catalyze the chemical modification of relevant metabolites. We previously showed that amyloids assembled with synthetic small peptides containing acidic residues can catalyze the hydrolysis of nucleotides, including adenosine triphosphate (ATP)^33,34^. α−Syn has a highly acidic 45-residues C-terminus preceded by a hydrophobic amyloid-prone middle region that can interact with nucleotides, including ATP. Thus, we tested whether purified recombinant α−Syn assembled into amyloids can chemically modify ATP and other nucleotides.

## Results

### α−Syn amyloids degrade ATP in a time-dependent manner

Purified α−Syn was seeded with previously assembled α−Syn amyloids to speed up aggregation. Upon incubation, the seeded amyloids were washed twice to remove monomers and potential contaminants. These isolated amyloids exhibited the characteristic Thioflavin-T fluorescence signal, with maximum emission centered around 488 nm, and classical amyloid fibril morphology by TEM analysis (supplementary figure 1). The isolated amyloids were then tested for activity. Initial trials showed phosphate release upon incubation of amyloids with ATP for 24 h at 37ºC in a Tris-buffer saline solution at pH 7.4. Thus, to get better insights on the products being formed we analyzed the reactions by HPLC, which was calibrated using nucleotide standards (Figure 1A). Interestingly, the reaction produced four major peaks associated to ATP, ADP, AMP and an unknown hydrolytic product. As expected, ATP remained mostly intact in a control reaction that lacked α−Syn amyloids. We then repeated the reaction in presence of ATP for different reaction times (Figure 1B). In presence of α−Syn amyloids and compared to time 0, ATP is highly degraded to all other forms at 24 h, reaching almost full degradation at 48 h. ATP remained almost intact even upon 48 h incubation in absence of amyloids.

**Figure 1.**
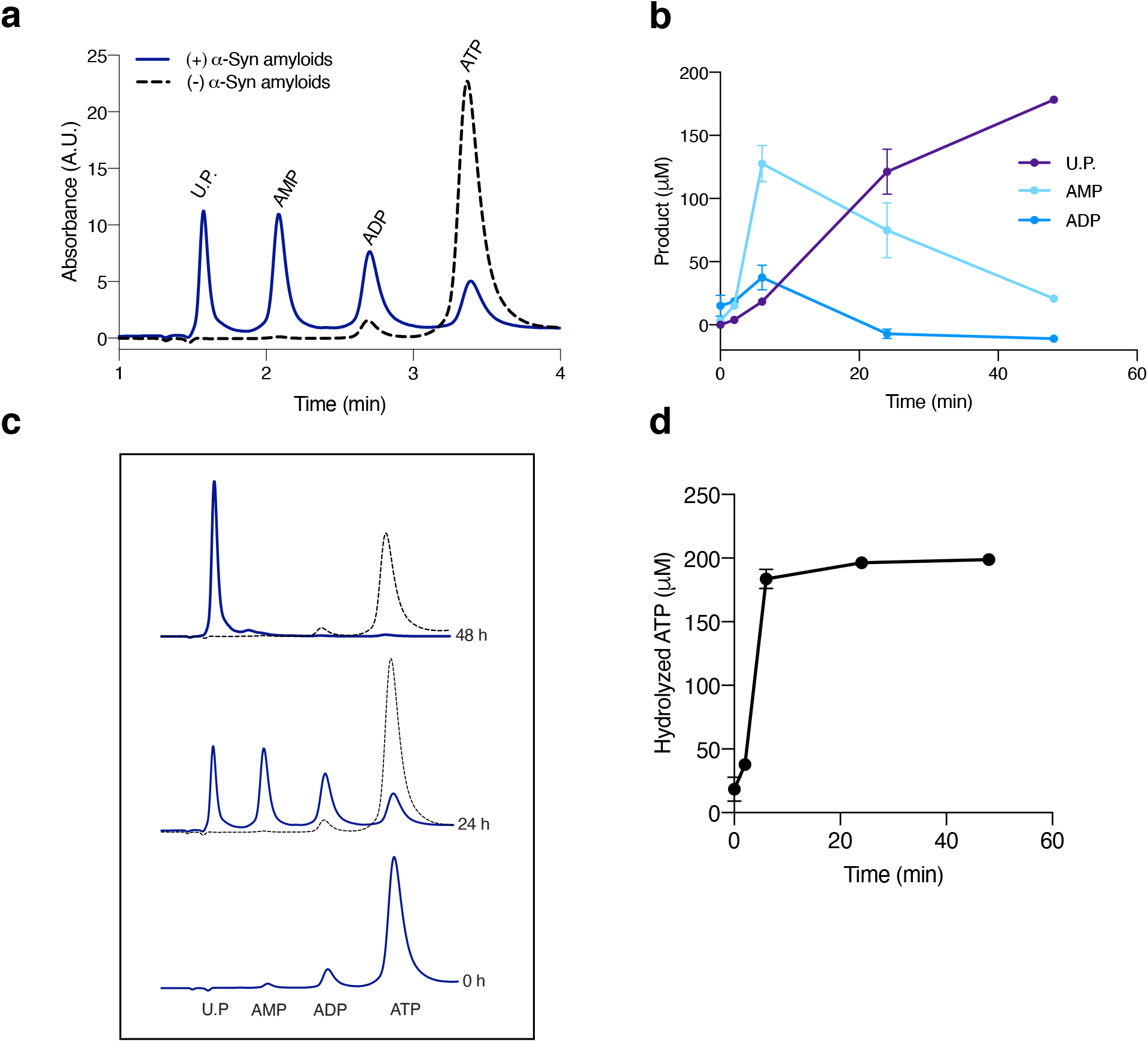
ATP degradation by α−Syn amyloids. (**a**) Products released from reactions run in triplicate for 24 h at 37ºC supplemented with 200 μM ATP in presence (straight dark blue line) or absence (dotted black line) of isolated α−Syn amyloids were analyzed by HPLC Absorbance (260 nm) was plotted as a function of the retention time with the indicated ATP degradation products. (**b**) The same reactions and controls were run at 0 h, 24 h and 48 h and analyzed by HPLC in the same fashion. (**c and d**). Evolution of the reaction products (**c**) from ATP degradation by α−Syn amyloids and hydrolyzed ATP (**d**) is plotted at different time points (0 h, 2 h, 6 h, 24 h and 72 h). Concentrations were estimated by the area of each peak. U.P. is unknown product.

Reactions at five different time points were run and analyzed by HPLC based on peak areas and the initial ATP concentration (Figure 1C). At 2 h, ADP and AMP had similar but low concentration levels whereas the unknown product was barely detectable. ADP and AMP reached maximum values at around 6 h whereas the unknown product peaked at 48 h. AMP maximum concentration at 6 h was around three times higher than that of ADP. At 24 h, ADP was already undetectable whereas AMP started decreasing until reaching one ninth of unknown product concentration at 48 h. The results agree with the estimated concentrations of hydrolyzed ATP (Figure 1D). At 2 h, ATP had already reached around 20% degradation. At 6 h around 90% of ATP was degraded, which increased to 98% at 24 h and to 99% at 48 h.

### ATP at physiological concentrations and AMP are substrates of α−Syn amyloids

ATP is present at very high concentrations in the cytoplasm of most cells. In neurons, ATP typically reaches cytoplasmic concentrations around 2 mM^35^. Therefore, we incubated isolated α−Syn amyloids with 2 mM ATP and analyzed the products by HPLC (Figure 2A). At 24 h, ATP degradation reached around 15%, with ADP and AMP having the highest levels. ADP reached a concentration 2.5 times greater than AMP whereas the unknown product concentration remained detectable but very low. A 72-h long reaction produced an ATP degradation of around 57%, with ADP again peaking, followed by AMP and the unknown product. Since AMP is the minimal nucleotide form derived from ATP, we hypothesized that the unknown product was being released by a different chemical reaction than hydrolysis of the phosphoanhydride bonds of ATP or ADP. Thus, we explored whether the unknown product can be directly released from AMP. Upon a 24-h incubation with Syn amyloids, the unknown product was clearly detected by HPLC whereas in absence of α−Syn amyloids the substrate remained intact. Hydrolyzed AMP reached around 60% and the unknown product had the same mobility and spectral property than the one found in the previous experiments.

**Figure 2.**
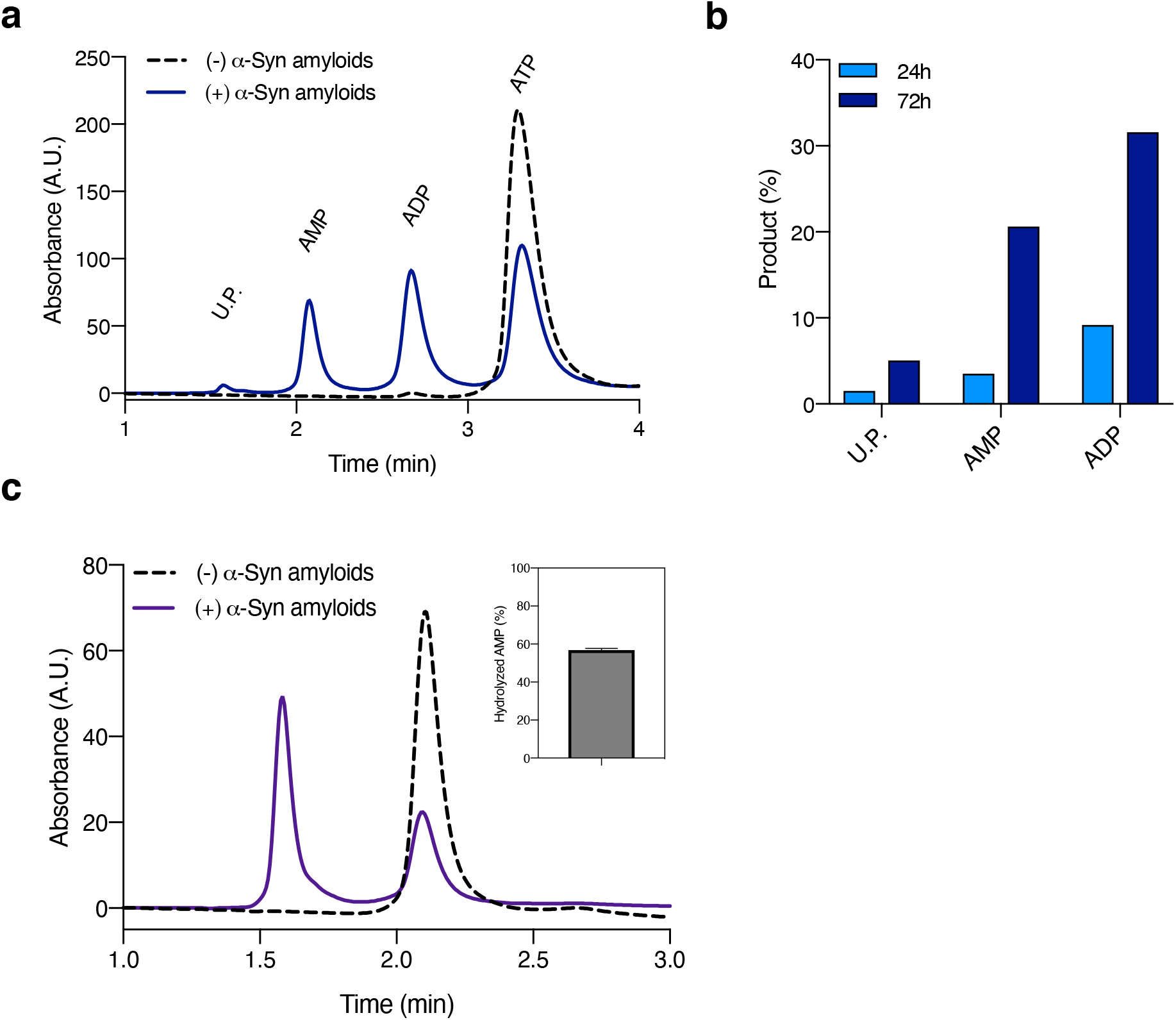
Degradation by α−Syn amyloids of AMP and physiological ATP. (**a**) Elution profile by HPLC of the products released from reactions run in triplicate for 24 h at 37ºC supplemented with 2 mM ATP in presence (straight dark blue line) or absence (dotted black line) of isolated α−Syn amyloids. (**b**) The same reaction in (a) was plotted along a 72 h-long reaction. Product concentration and percentage were estimated by the area of each peak. (**c**) Elution profile of AMP (200 μM) incubated with isolated α−Syn amyloids for 24 h at 37ºC. The inset shows the amount of hydrolyzed ATP quantified from the peak areas. U.P. is unknown product.

Since ATP degradation proceeded exclusively in the presence of α−Syn amyloids, it is likely that the degradation products are released in a catalytic fashion^33^. Thus, we analyzed whether degradation followed a catalytic enzyme-like behavior. We mixed α−Syn amyloids with ATP at different concentrations for 2 h, a time interval in which the degradation reaction appears linear with time (Figure 1C and 1D). All the products were quantified by HPLC as previously mentioned and the data was fitted to a Michaelis-Menten model. The results show that α−Syn amyloids degrade ATP through a classical substrate saturation curve (Figure 3). The fitted value for *K*_*M*_ was around 200 μM (the concentration we used in most of the previous experiments).

**Figure 3.**
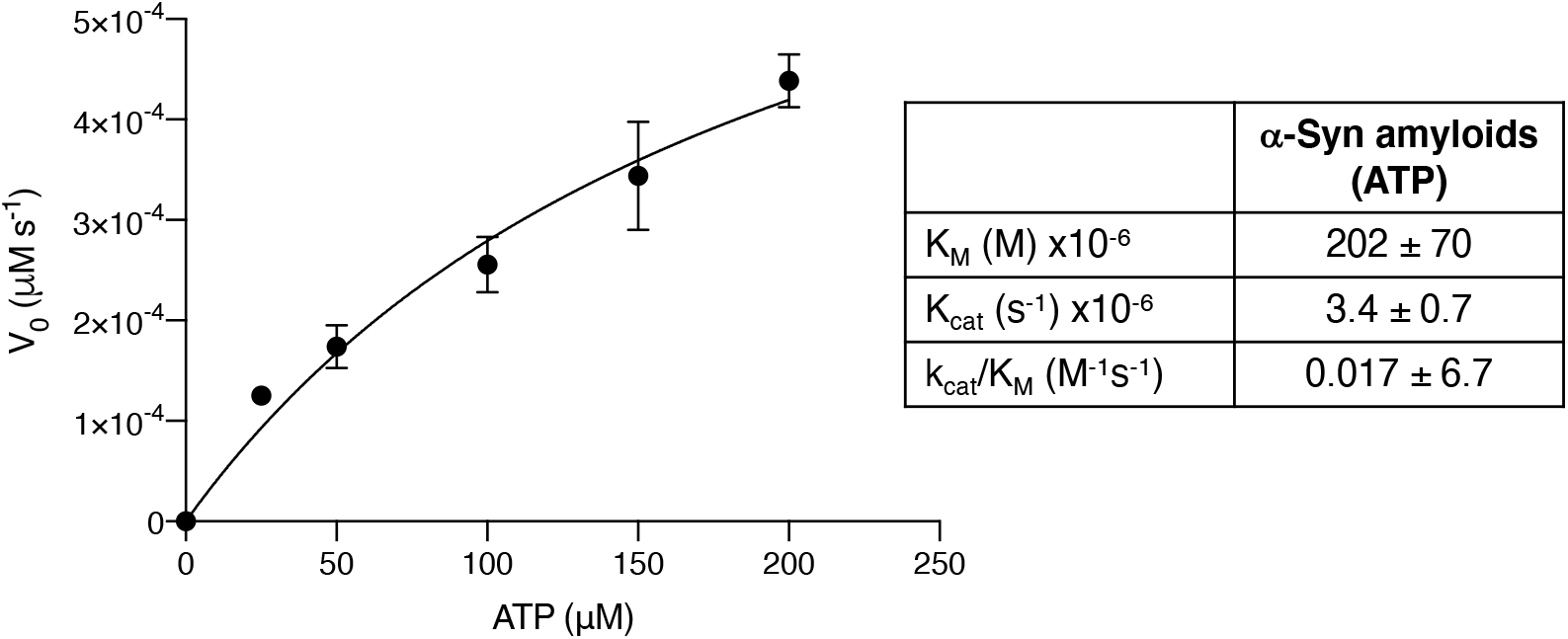
Substrate saturation curve of ATP degradation by α−Syn amyloids. Isolated α−Syn amyloids were incubated in TBS with different ATP concentrations in triplicate for 2 h at 37ºC. The products were analyzed by HPLC, and the areas of all ATP degradation products were counted as total product concentration. The data was fitted through nonlinear fit using a Michaelis-Menten model. The kinetic parameters: Michaelis constant, *K*_*M*_, catalytic constant, *k*_*cat*_, and catalytic efficiency, *k*_*cat*_/*K*_*M*_ are shown in the table.

**Figure 4.**
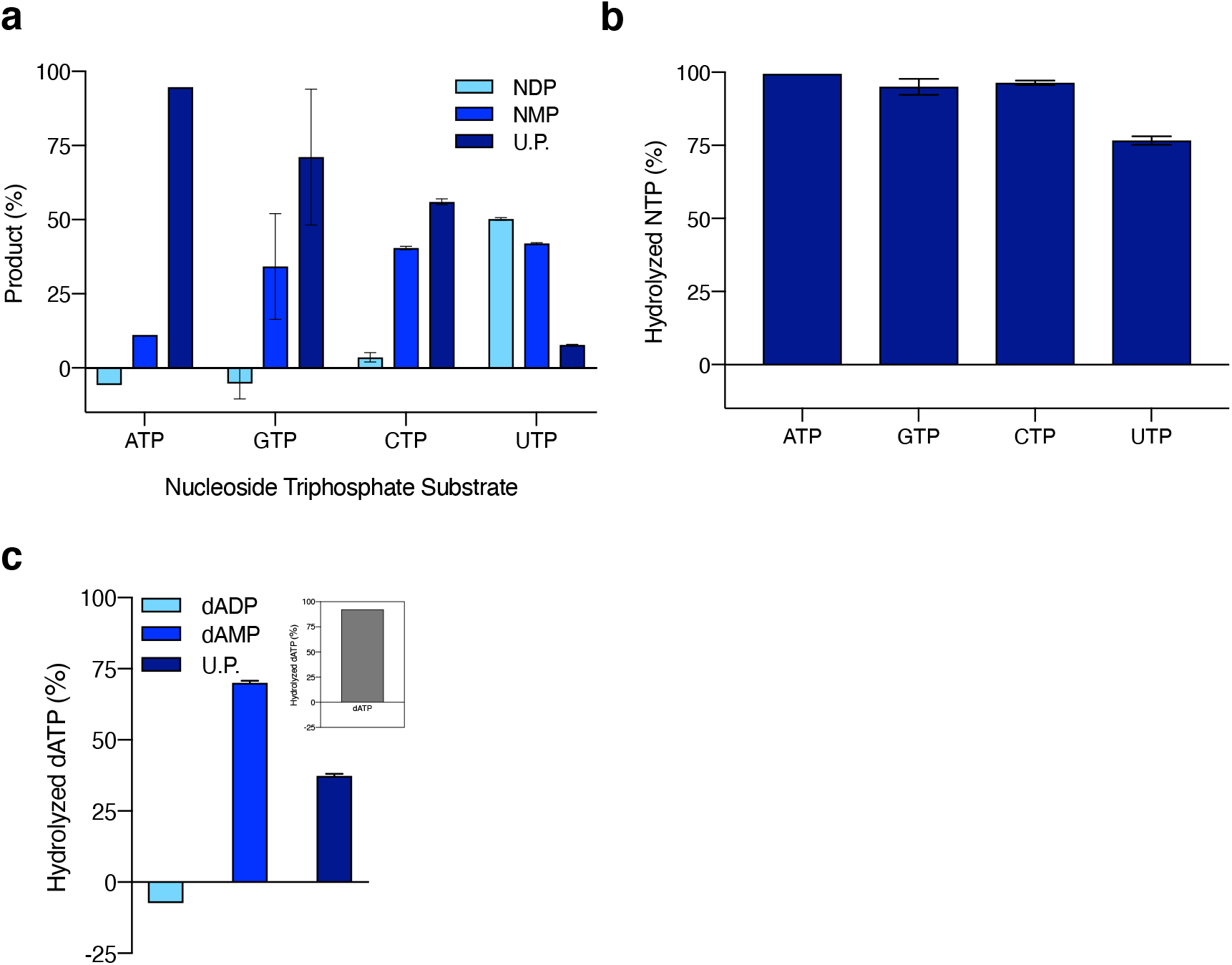
Degradation by α−Syn amyloids of other nucleotides. (**a**) Concentrations (expressed as percentage) of all degradation products determined by HPLC from reactions run in triplicate for 24 h at 37ºC supplemented with 200 μM of different nucleotides (ATP, GTP, CTP or UTP) in presence (straight dark blue line) or absence (dotted black line) of isolated α−Syn. (**b**) Hydrolyzed amount of each ribonucleotide expressed as percentage. (**c**) Concentrations of degradation products from a reaction containing isolated α−Syn amyloids supplemented with 200 μM dATP for 24 h at 37ºC. The inset shows the hydrolyzed dATP expressed as percentage. U.P. is unknown product.

### α−Syn amyloids can degrade other nucleotides

We explored whether α−Syn amyloids can degrade other ribonucleotides, specifically guanosine triphosphate (GTP), cytidine triphosphate (CTP) and uridine triphosphate (UTP). We incubated isolated α−Syn amyloids with each ribonucleotide in separate tubes for 48 h and analyzed the products by HPLC. The results showed that α−Syn amyloids can effectively degrade all the ribonucleotides albeit with different efficiencies. All three nucleoside triphosphates were degraded to nucleoside di- and monophosphates. As with ATP, we also observed the formation of a third hydrolytic product of unknown nature. Compared to ATP, GTP was the second most effectively degraded ribonucleotide, followed by CTP and UTP. ATP, GTP and CTP showed minor accumulation of nucleoside diphosphate, and the dominating species was the unknown product. For UTP, the reaction appear less efficient since uridine diphosphate is the main product, followed by uridine monophosphate and an unknown species. GTP and CTP reached almost full depletion (>90%) whereas UTP remained at around 75%. We also tested whether other physiologically relevant nucleotides can be degraded by α−Syn amyloids, such as deoxyadenosine triphosphate (dATP) as the deoxyribonucleotide ATP counterpart, and a cyclic derivative of AMP (3’-5’ cAMP), a second messenger essential in signal transduction. A 48-h incubation of α−Syn amyloids and dATP led to almost full degradation (92%) of the nucleotide, releasing dAMP as the main species followed by an unknown product. As observed with ATP degradation, dADP was below the detection level. On the other hand, α−Syn amyloids were unable to degrade 3’-5’ cAMP, which remained virtually unaltered upon 48-h incubation.

## Discussion

Our results evidence a novel gain-of-function for α−Syn amyloids that can have direct implications in pathological ATP metabolism. The degradation releases all hydrolytic nucleoside phosphate derivatives. ATP is rapidly degraded, and the remaining species are progressively degraded into subsequent products. The degradation is clearly catalytic and enzyme-like with substrate saturation. Although the present data is insufficient to determine a mechanistic pathway, the results provide strong evidence for a sequential degradation ATP -> ADP -> AMP -> unknown product, as these products accumulate orderly in time. This is further supported by the degradation acting directly over AMP and apparently yielding the same unknown product. Although we initially suspected this product to be adenosine, produced by hydrolysis of the remaining phosphate of AMP, its HPLC mobility and spectral properties indicate a different nature. We are currently studying this and the mechanistic aspects.

Interestingly, ATP was degraded even at high and physiological concentrations. This is intriguing since ATP is currently recognized as a biological hydrotrope, meaning it can avoid protein aggregation by increasing the solubility of different proteins, including those prone to form amyloids^36^. However, recent findings demonstrated that the hydrotropic effects on α−Syn are complex, with a biphasic behavior in which low ATP concentrations reduce the aggregation lag-phase whereas high concentrations reduce the aggregation plateau^37^. The latter occurs only at very high and non-physiological ATP concentrations (> 10 mM), which can explain why α−Syn amyloids such as Lewy bodies can form and endure in the cytoplasm, especially in neurons in which ATP is around 2 mM. In fact, 2 mM ATP accelerates α−Syn aggregation without modifying the plateau. Our kinetic data show that at 2 mM, the activity of α−Syn amyloids is well in the saturated region, implying the amyloids are virtually at full catalytic operating capacity.

The association of α−Syn and ATP goes beyond hydrotropic effects^38^. ATP naturally binds *in vitro* to α−Syn monomers through electrostatic and nonpolar interactions, and since ATP is abundant in cells, such interactions may be ubiquitous and frequent in the cytoplasm^37^. In fact, we found in preliminary experiments that several nucleotides co-purify with α−Syn, which are effectively removed in the washing steps. It is thus plausible that ATP may not only colocalize with α−Syn amyloids inside cells but also accelerate and stabilize their formation at physiological concentrations, while promoting its own degradation.

There is abundant literature reporting impaired bioenergetics in neurodegenerative diseases, including PD^38–40^. Energy-demanding synapses and cytoplasmatic ATP levels are reduced early in PD, preceding neuronal death^41^. Dopaminergic neurons are particularly sensible to ATP levels due to constant dopamine production and release, two highly ATP-consuming processes^42,43^. Energy-impairment is partially associated to mitochondrial dysfunction in PD. α−Syn amyloids can bind and disrupt mitochondrial membranes and impair respiratory ATP production while increase ROS^20–23^. However, energy deprivation can also be linked to cytoplasmatic factors. Activation of phosphoglycerate kinase 1 (the first ATP-producing enzyme in glycolysis) by terazosin has shown significant protection against neurodegeneration and cell death in cellular and animal models of PD, and in several clinical studies^44,45^. Indeed, terazosin was recently reported to effectively boost ATP levels, suppressing metabolic stress in dopaminergic neurons, and acting as a leverage of metabolic impairments in PD^46^.

All this evidence points that optimal ATP levels are crucial in PD. Mitochondrial and cytoplasmatic ATP metabolism are evidently interconnected and thus diverse PD-associated metabolic dysfunctions are inescapably complementary. Excess ROS production and low ATP levels may act concertedly in pathogenesis. For instance, low cytoplasmatic ATP levels triggered by α−Syn amyloids may produce an initial hyperactive mitochondrial state, which can also produce ROS. Mitochondrial hyperactivity is an early pathological marker in worm models of PD, which led to propose decreased mitochondrial function (typically observed in postmortem analysis) as a possible end stage in PD^47,48^. Future studies on the cellular implications of ATP degradation by α−Syn amyloids may shed new light on metabolic disfunction in PD. Overall, our results may underly a previously unrecognized pathogenic mechanism that could redefine ATP metabolism in PD and other synucleinopathies, paving the way for future targeted therapeutic interventions.

## Materials and Methods

### Protein expression and purification

Recombinant α−Syn was expressed in *E. coli* (DE3) cells, which were grown at 37ºC under agitation in LB medium supplemented with ampicillin and induced with IPTG at O.D. around 0.6. Cells were harvested by centrifugation at 14,000 x g and were suspended in a solution containing 25 mM Tris-HCl pH 7.8 and 750 mM NaCl. The suspension was boiled for 20 min and centrifuged at 18,000 x g for 30 min and the pellet discarded. The supernatant was dialyzed against 25 mM Tris-HCl pH 7.8 and 50 mM NaCl overnight (dialysis membrane of 3500 MWCO), and then loaded on a 5 mL HiTrap Q HP column (Cytiva) and eluted with a linear gradient (50-500 mM NaCl). The protein-containing fractions were collected, pooled together, and dialyzed against a solution with 10 mM Tris-HCl pH 7.8 and 25 mM NaCl. Upon dialysis, the sample was centrifuged at 20,000 x g for 30 min and the soluble fraction was syringe-filtered (0.22 μM). The protein was concentrated using a 10,000 MWCO Amicon centrifugation filter (Milipore). Purity was confirmed by SDS-PAGE and the protein was separated in single-use aliquots and kept frozen at -40ºC.Final protein concentration was estimated using a standard BCA protocol (Pierce).

### In vitro α−Syn aggregation

Amyloid formation was induced by addition of diluted premade α−Syn amyloids (seeds) to α−Syn monomers. The seeds were prepared by incubating freshly thawed α−Syn monomers at 250 μM in TBS buffer (50 mM Tris-HCl pH 7.4 and 150 m NaCl) for 8 days at 37ºC with continuous agitation. Upon completion, the seeds solution was stored frozen at -40ºC. The formation of α−Syn amyloids for the catalytic reactions was produced by mixing freshly thawed α−Syn seeds with monomer α−Syn, yielding a final concentration of 250 μM in TBS solution. The reaction was incubated for 5 days at 37ºC with continuous agitation. To separate amyloids from monomers and potential contaminants, the resulting α−Syn amyloids were washed by centrifugation for 30 min at 31,000 x g, the supernatant was discarded, and the pellet resuspended in TBS solution. This washing step was repeated, and the final pellet (isolated α−Syn amyloids) was resuspended in TBS or solution for catalytic reaction.

### TEM analysis and Thioflavin-T assay

Samples containing isolated α−Syn amyloids were analyzed by transmission electron microscopy (TEM) and Thioflavin-T assay. For Thioflavin-T assay, isolated α−Syn amyloids in TBS were supplemented with 25 μM ThT and the reaction was analyzed in triplicate by emission fluorescence in a plate reader (Tecan M200 Pro Infinity) with excitation set at 435 nm. For TEM analysis, 10 μL of isolated α−Syn amyloids in TBS were deposited for 5 min on previously ultraviolet light-activated carbon grids. The grids were then soaked in milli-Q water and negative stained with 2% uranyl acetate for 2 min, followed by a final soaking step. The grids were analyzed in a Talos F200 G2 (ThermoFisher Scientific) transmission electron microscope, loaded with a Ceta 16M CMOS camera (16 bit) and the images were acquired and processed with a Velox imaging software.

### ATP degradation assay

Isolated α−Syn amyloids (200 μM) were suspended in TBS solution supplemented with different concentrations of ATP or other nucleotides. Control reactions lacking α−Syn were treated and run in parallel. The samples were incubated at 37ºC for different times (specified in the corresponding figures) under continuous agitation. The reaction products were analyzed by high performance liquid chromatography (HPLC) or with a colorimetric kit for phosphate determination. For HPLC, the samples were syringe-filtered through a 0.22 μM filter and analyzed by HPLC (Agilent Infinity 1220 system loaded with diode-array detector) using an ion-pair reverse chromatography, as reported previously. Briefly, the samples were automatically loaded on a C18 column previously equilibrated with a mobile phase containing 50 mM K_2_HPO_4_, 20 mM glacial acetic acid and 3-7% (V/V) acetonitrile and followed at 260 nm. Commercial nucleotides were used as standards under the same conditions to determine elution times, which remained stable throughout all the experiments. Product concentrations were determined using the in-house software by estimating the area under the curve of all the peaks and weighted by the total nucleotide concentration. For the colorimetric assay, the total phosphate content was estimated by using a commercial kit based on the malaquite-green assay and following the kit instructions (Enzo). The samples were spectrophotometrically analyzed in a plate reader set at 620 nm. All the results were analyzed and plotted using Prism Software (Graphpad).

## Author Contributions

C.CC. and R.DE. conceived the idea. C.CC. and E.N. performed the experiments. C.CC. and R.DE. analyzed the data. R.DE. wrote the manuscript.

## Funding Support

This research was funded by Chilean research grant ANID FONDECYT 13220108

## Data availability statement

All data generated and analyzed during this study are included in this published article and its Supplementary Information.

## Additional information

Supplementary information accompanies this paper.

## Competing Interests

The authors declare no competing interests.

## Notes

### Competing Interest Statement

The authors have declared no competing interest.

