## Supplemental figure for "Alpha-synuclein amyloids catalyze the degradation of ATP and other nucleotides"

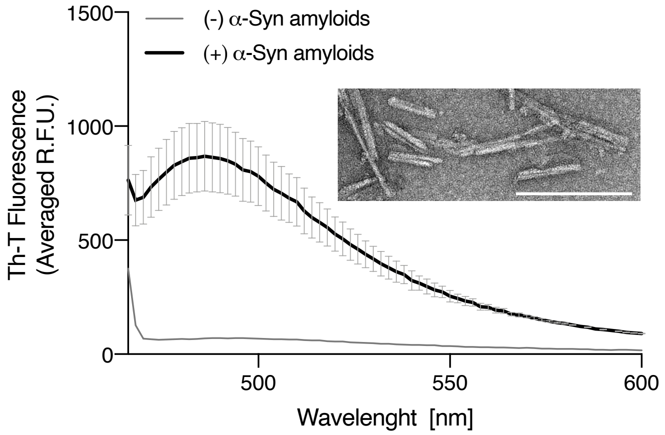


**Figure S1**. Characterization of the amyloid state of α–Syn. Fluorescence emission spectrum of isolated α–Syn fractions in presence of Thioflavin-T (25 μM) with excitation set at 435 nm. The inset shows a micrograph of isolated α–Syn amyloids analyzed by TEM (white bar indicates 200 nm). U.P. is unknown product.
